# Phenotyping maize seed tolerance to storage after seed treatment using a Seed Treatment Tolerance Index

**DOI:** 10.64898/2026.03.09.710582

**Authors:** Venicius U. V. Reis, Giulyana I. S. Tavares, Danilo C. Maciel, Jaqueline P. Januário, Marina S. R. Pereira, Raquel M. O. Pires, Everson R. Carvalho

**Affiliations:** Department of Crop Sciences, Federal University of Lavras, Lavras, MG, Brazil; Department of Plant Sciences, University of California, Davis, CA, USA; Agriculture, Food Systems & Bioeconomy Research Centre, Ryan Institute, University of Galway, Galway, Ireland

**Keywords:** *Zea mays* L., seed quality, neonicotinoids, multivariate phenotyping, phytotoxicity, abiotic stress

## Abstract

Neonicotinoid seed treatments protect maize during early growth but can induce phytotoxicity that intensifies during storage. Despite recognized genotypic variation in tolerance, standardized phenotyping methods are lacking. We evaluated nine commercial maize hybrids under three seed treatments (control, one neonicotinoid [1N], and two neonicotinoids [2N]) across two storage periods (0 and 6 months at 25 °C) using germination, accelerated aging, and cold tests. A Seed Treatment Tolerance Index (STTI) was analyzed through hierarchical clustering, principal component analysis, and multivariate analysis of variance. Results showed a significant triple interaction among genotype, seed treatment, and storage. Hybrids from female line A maintained STTI above 0.95, while female C hybrids showed germination reductions up to 48 percentage points and vigor losses up to 90 percentage points under 2N after six months. Tolerance was associated with hydrogen peroxide regulation by catalase and ascorbate peroxidase. The STTI proved a reliable tool for classifying genotypic tolerance, with direct applications for breeding programs and seed industry logistics.

## INTRODUCTION

High physiological quality of maize seeds is essential for successful agricultural production. Germination and early seedling vigor determine stand establishment and, consequently, final crop yield (Reis et al., 2022; Reed et al., 2022). Technologies that protect seeds during their early stages are therefore widely adopted. Among these, seed treatment (ST) with phytosanitary products has consolidated itself as a near-universal practice in agriculture, with the maize ST market valued at USD 3.32 billion (Reis et al., 2026a). Its widespread adoption is justified by the effective protection it provides to seeds and seedlings against early pests and diseases (Neves et al., 2022; Reis et al., 2026a).

However, this technology carries a significant risk of phytotoxicity, which can induce negative physiological responses and compromise seed quality. Despite this risk, neonicotinoids remain an essential component of maize ST, providing systemic protection against key pests such as the *Spodoptera frugiperda*, with residual efficacy extending up to 35 days after emergence and documented yield reductions above 40% when omitted (Pes et al., 2020; Ismail, 2024). This problem is particularly well documented with neonicotinoid insecticides, which represent the most phytotoxic insecticide class used in ST (Tamindžić et al., 2016; Shahid et al., 2024). In this context, phytotoxicity refers to the cellular damage induced by the chemical agent (Taylor and Salanenka, 2012). This damage is frequently aggravated by storage, which compounds the natural seed deterioration process (Moraes et al., 2022; Reis et al., 2026a).

The underlying mechanism of this damage is linked to biochemical disturbances, primarily the induction of oxidative stress. The overproduction of reactive oxygen species (ROS) can lead to lipid peroxidation, compromising membrane integrity and the function of vital organelles such as mitochondria, thereby affecting the energy required for germination (Shahid et al., 2021; Zhang et al., 2022; Forti et al., 2024). A genotype’s capacity to tolerate this stress depends on the efficiency of its antioxidant defense system. Despite recognition of this genotypic variability, a standardized phenotyping methodology for its robust evaluation is still lacking (Tamindžić et al., 2016; Reis et al., 2026b).

Current assessments, often based on a single physiological test (Reis et al., 2026b), are insufficient to capture the complex interaction among genotype, treatment, and storage. This limitation hinders precise genotype classification, representing a bottleneck for breeding programs and decision-making at the time of ST. An integrated approach that combines multiple tests into a tolerance index is therefore necessary.

Given the consistent genotypic differences in tolerance to seed treatment and storage, and the evidence that these can be identified through a weighted combination of physiological quality tests and antioxidant system responses, the main objectives of this study were: (i) to quantify tolerance among maize genotypes to the synergistic stress of neonicotinoid seed treatment and storage, (ii) to elucidate the biochemical mechanisms associated with tolerance, focusing on the antioxidant defense system, and (iii) to develop and validate a multivariate methodology for classifying these genotypes based on a Seed Treatment Tolerance Index (STTI).

## MATERIALS AND METHODS

### Plant Material and Experimental Design

Seeds from nine commercial maize hybrids and their parental lines, harvested in the 2022/2023 season, were evaluated in a completely randomized design with a 9 × 3 × 2 factorial arrangement (9 genotypes × 3 seed treatments × 2 storage periods), with four replications of 50 seeds per unit. Genotype identities and parental crosses are described in Table S1 of the Supporting Information.

### Seed Treatments and Storage Conditions

Seeds were treated in 2-kg batches using an industrial-batch simulator (Arktos Laboratório L2K-BM, BM Momesso, Matão, Brazil), calibrated at 15 Hz to simulate industrial batch-type treatment. Three treatments were applied (Table S2): control (fungicide mixture plus polymer), 1N (control plus one neonicotinoid plus one diamide insecticide), and 2N (control plus two neonicotinoid insecticides). After treatment, seeds were stored for six months in a B.O.D. chamber (25 ± 2 °C) to simulate uncontrolled storage in tropical environments, with evaluations performed at 0 and 6 months.

### Physiological Quality and Phytotoxicity

Physiological quality was assessed through four tests: rolled paper germination (RP), rolled paper + vermiculite (RP+V), accelerated aging (AA), and cold test (CT), (Cicero and Vieira, 2020; Marcos-Filho, 2020; Rocha et al., 2023; Brasil, 2025). From these tests, the phytotoxicity index (Pi) was calculated for each variable as the relative performance reduction of insecticide treatments relative to the control (Reis et al., 2026b). Complete protocols are described in Methods S1 of the Supporting Information.

### Seed Treatment Tolerance Index

The seed treatment tolerance index (STTI) was calculated for each combination of genotype, treatment, and storage period across all four physiological variables, as the ratio of mean performance under insecticide treatment to the control mean (Equation 1):

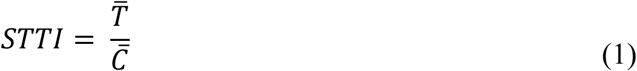

where T is the mean of the analyzed variable under the insecticide treatment (1N or 2N) and C is the mean under the control for the same genotype and storage period. Values near 1.0 indicate full tolerance; values near 0 indicate complete sensitivity. A general STTI was obtained as the arithmetic mean across all physiological variables and both treatments (1N and 2N), yielding one summary index per genotype. This normalization minimizes the influence of intrinsic genotypic deterioration differences, enabling direct comparison of treatment impact across genotypes.

### Biochemical Analyses

The enzymatic activities of superoxide dismutase (SOD), catalase (CAT), and ascorbate peroxidase (APX), and hydrogen peroxide (H₂O₂) content (Giannopolitis and Ries, 1977; Nakano and Asada, 1981; Havir and McHale, 1987; Azevedo et al., 1998; Velikova et al., 2000), were measured in the two most contrasting hybrids. A 2 × 3 × 2 factorial arrangement was used with three replications of 200 mg of macerated seeds. Complete protocols are in Methods S1 of the Supporting Information.

### Statistical Analysis

Individual ANOVAs were performed for each variable, with means compared by the Scott-Knott test (α ≤ 0.05) using the ExpDes.pt package (Ferreira et al., 2021). Genotype classification was performed by sequential multivariate analysis using Pi and STTI as input variables in the MVar.pt package (Ossani and Cirillo, 2025). Data were standardized (z-score) and submitted to hierarchical cluster analysis using Ward’s method with Euclidean distance, visualized by PCA. Cluster distinction was verified by MANOVA using Wilks’ Lambda followed by univariate ANOVAs. The final tolerance classification was based on the general STTI at six months of storage. All analyses were conducted in R studio (R Core Team, 2024).

## RESULTS

### Physiological Quality Assessment

In the RP test, at the beginning of storage, no significant difference in germination was observed among genotypes under the control seed treatment. Under 1N, genotype 8C showed significantly lower germination than the others. Under 2N, genotypes from female line C (6C, 7C, and 8C) recorded germination up to 12 percentage points (pp) lower than those from female line A (1A, 2A, and 3A) (Figure 1). After six months, differences were amplified: under 2N, the difference between the most tolerant genotype from female A (2A) and the most sensitive from female C (8C) reached 48 pp. Storage caused significant germination losses in sensitive genotypes under chemical treatment, with the greatest reductions under 2N (24 pp for 4B and 6C, and 36 pp for 8C). In contrast, genotypes from female A maintained high stability, with maximum losses of only 1 pp across all treatments.

**Figure 1.**
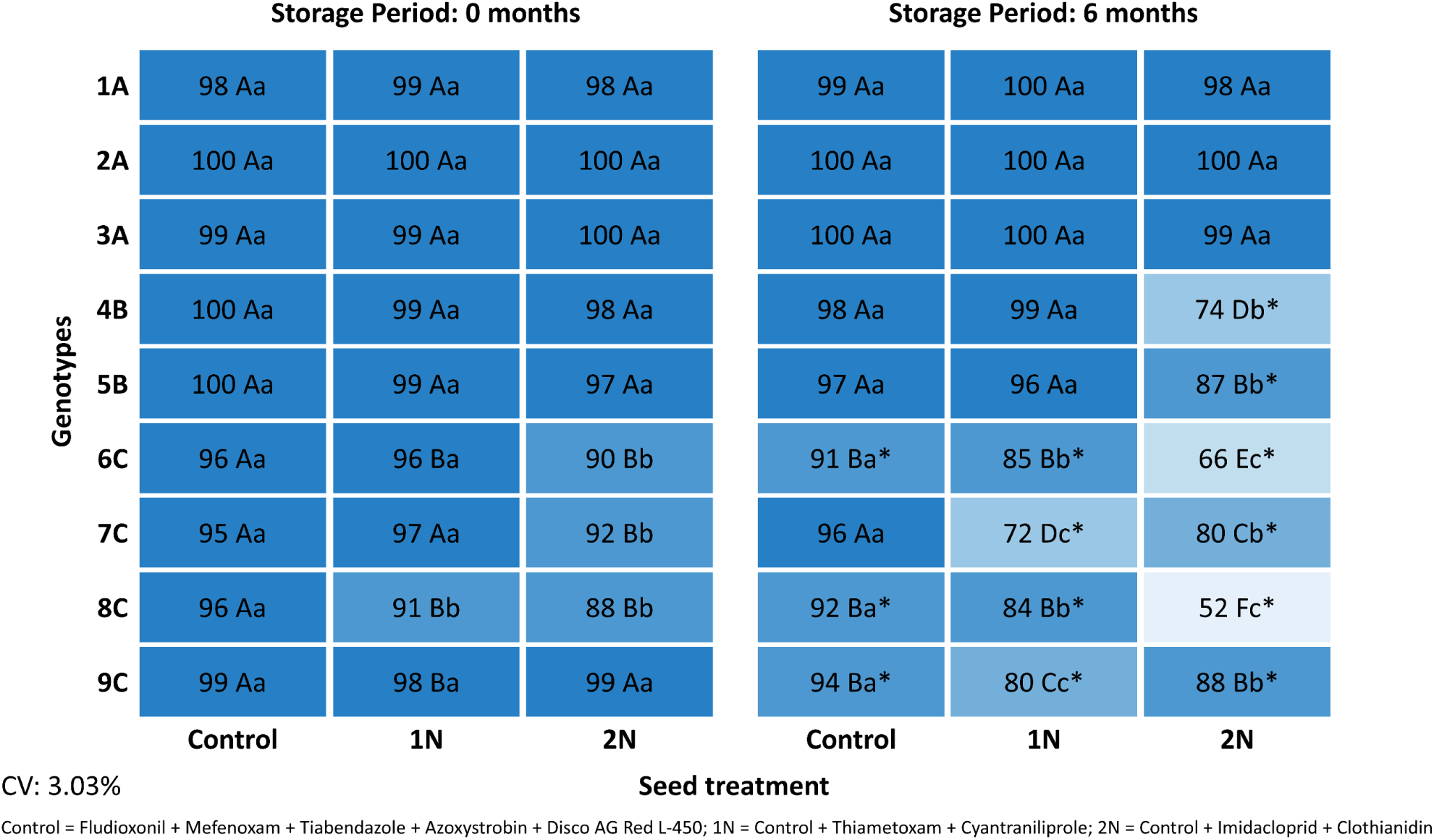
Percentage of normal seedlings of maize in the rolled paper germination test (7 days) as a function of genotype, seed treatment, and storage period (0 and 6 months at 25 °C). Seed treatments: Control (fungicide + polymer), 1N (control + one neonicotinoid + one diamide insecticide), and 2N (control + two neonicotinoid insecticides). Means followed by different uppercase letters differ among genotypes within the same column (treatment and storage period); different lowercase letters differ among seed treatments within the same row (genotype and storage period); asterisks (*) indicate significant differences between storage periods for the same genotype and treatment (Scott-Knott test, p ≤ 0.05).

In the RP+V test, differences among genotypes were not detected under the control, but under 2N, female C genotypes recorded germination up to 16 pp lower than female A genotypes at the beginning of storage (Figure 2). After six months, the performance hierarchy was maintained. Under 2N, germination of genotype 3A exceeded that of 8C by 23 pp. Storage reduced germination in most treatment combinations, with minimal effect on female A genotypes (reduction ≤ 3 pp) and a significant impact on female C under treatments 1N and 2N, and female B under 2N. For example, genotype 6C reached initial germination of 93% under 2N, declining to 78% after six months (15 pp).

**Figure 2.**
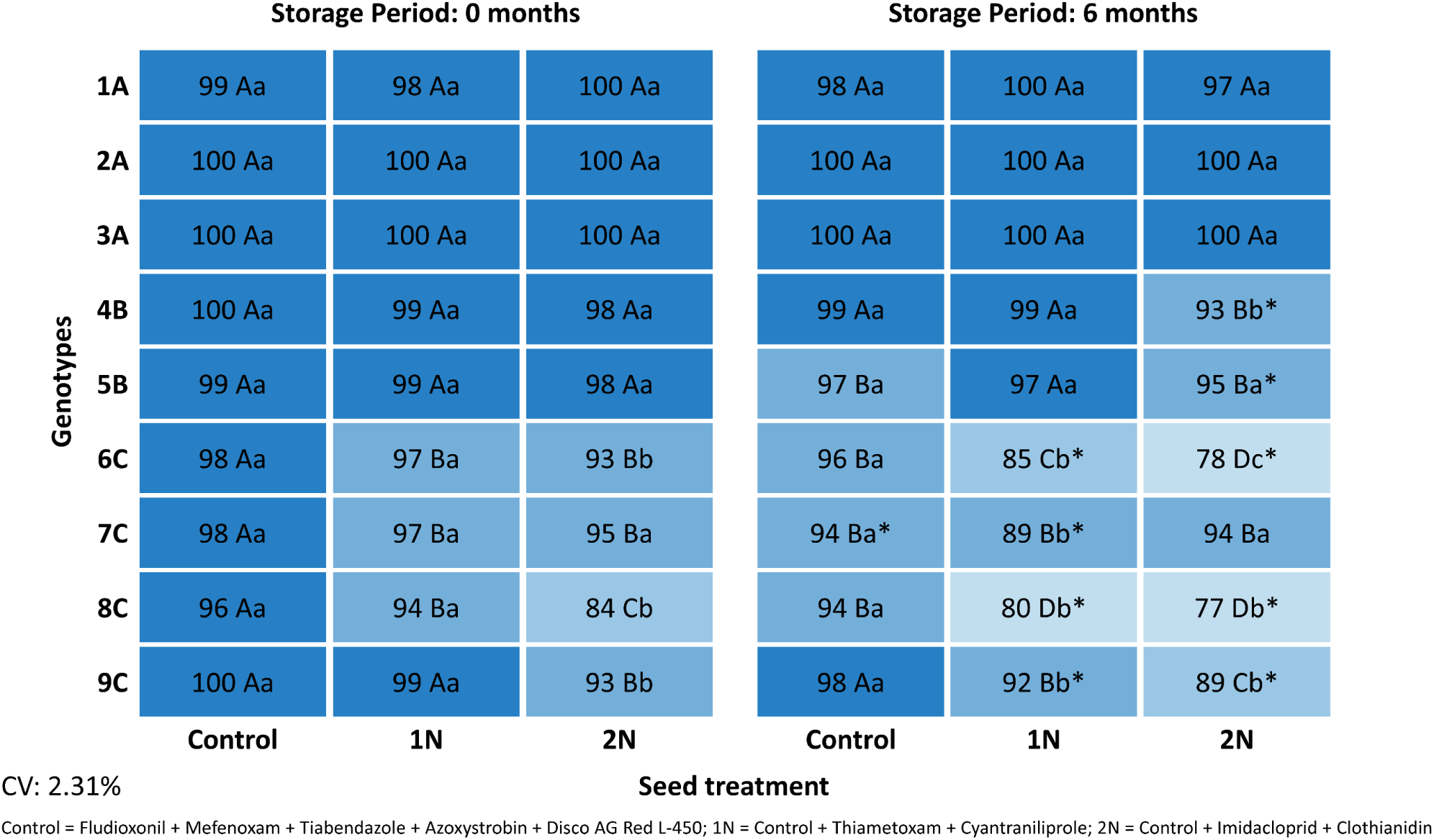
Percentage of normal seedlings of maize in the rolled paper plus vermiculite germination test (7 days) as a function of genotype, seed treatment, and storage period (0 and 6 months at 25 °C). Statistical comparisons and notation as described in Figure 1.

In the AA test, which subjects seeds to high temperature and humidity, differences among genotypes were even more pronounced (Figure 3). At the beginning of storage, even in the control, genotype 7C recorded lower vigor (77%) than genotypes from female lines A and B, followed by 8C and 9C. Under 2N, this initial difference for 7C increased to 55 pp relative to genotype 1A. After six months, vigor losses were amplified, especially in female lines B and C. In the control, 7C lost 68 pp, while 6C under 2N lost 45 pp. In contrast, female A genotypes showed high tolerance, with a maximum loss of 16 pp after six months (2A), even under 2N. Under 1N after storage, the vigor difference between female lines A and C exceeded 90 pp.

**Figure 3.**
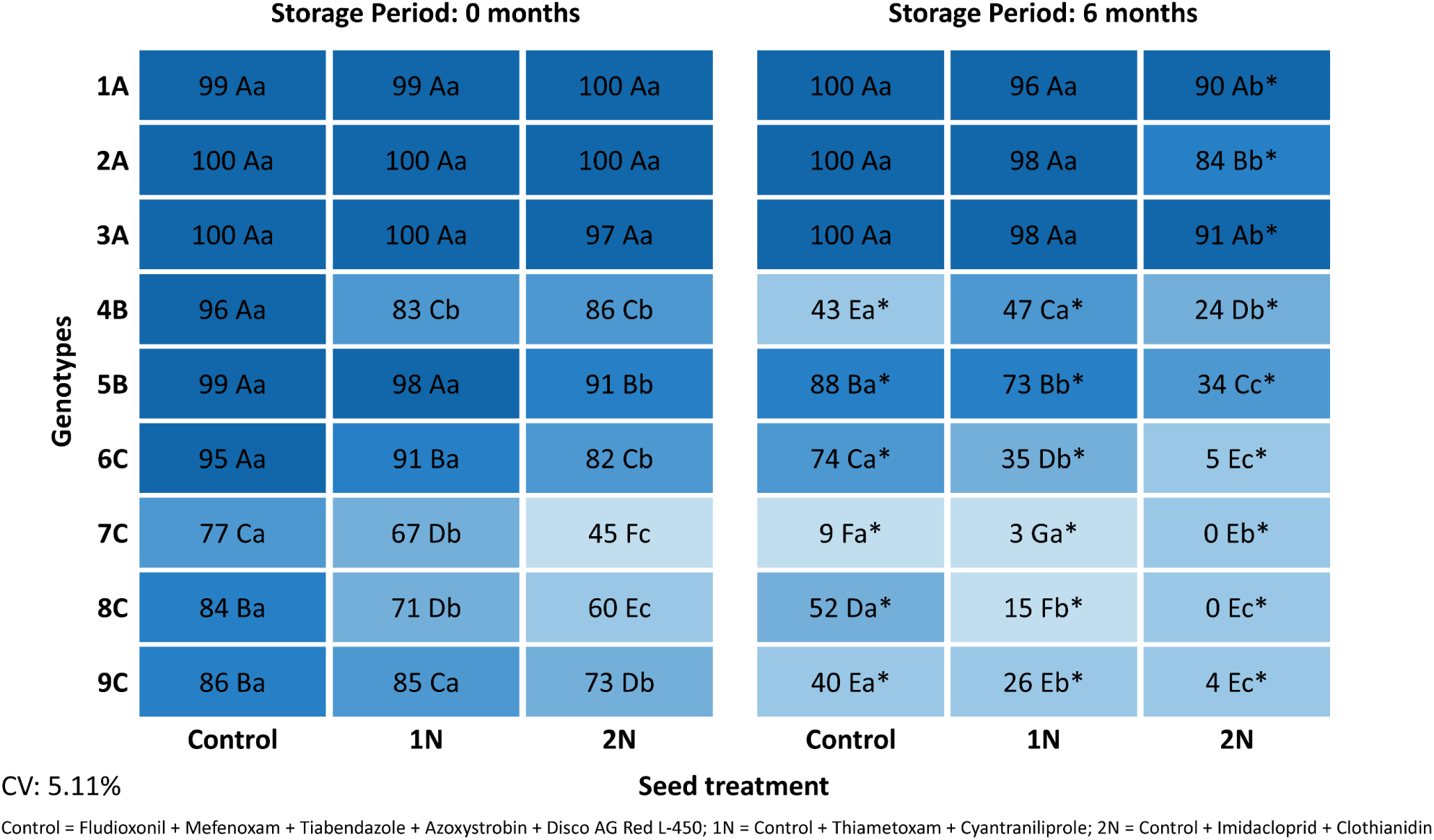
Percentage of normal seedlings of maize in the accelerated aging test (41 °C, 72 h), evaluated by the germination test at 4 days, as a function of genotype, seed treatment, and storage period (0 and 6 months at 25 °C). Statistical comparisons and notation as described in Figure 1.

In the CT test, which simulates low-temperature sowing conditions, differences among factors were also observed (Figure 4). At the beginning of storage, differences among genotypes were already visible in the control, with genotype 8C recording emergence 7 pp lower than the most vigorous genotypes, alongside 4B and 7C. The negative effects of the treatments were immediate (8C showed a 19 pp reduction in emergence under 2N relative to the control). After six months, the combination of storage and chemical treatment intensified differences. Under 2N, more tolerant genotypes such as 3A surpassed 8C by more than 51 pp, with 8C reaching only 49% emergence. Other genotypes, such as 4B and 7C, suffered reductions of 13 to 26 pp after storage with 2N. In contrast, some female A and B genotypes (1A, 3A, and 5B) maintained high emergence across all scenarios, with 3A preserving high emergence in every condition and 1A showing a minimal reduction of only 2 pp under 2N after six months.

**Figure 4.**
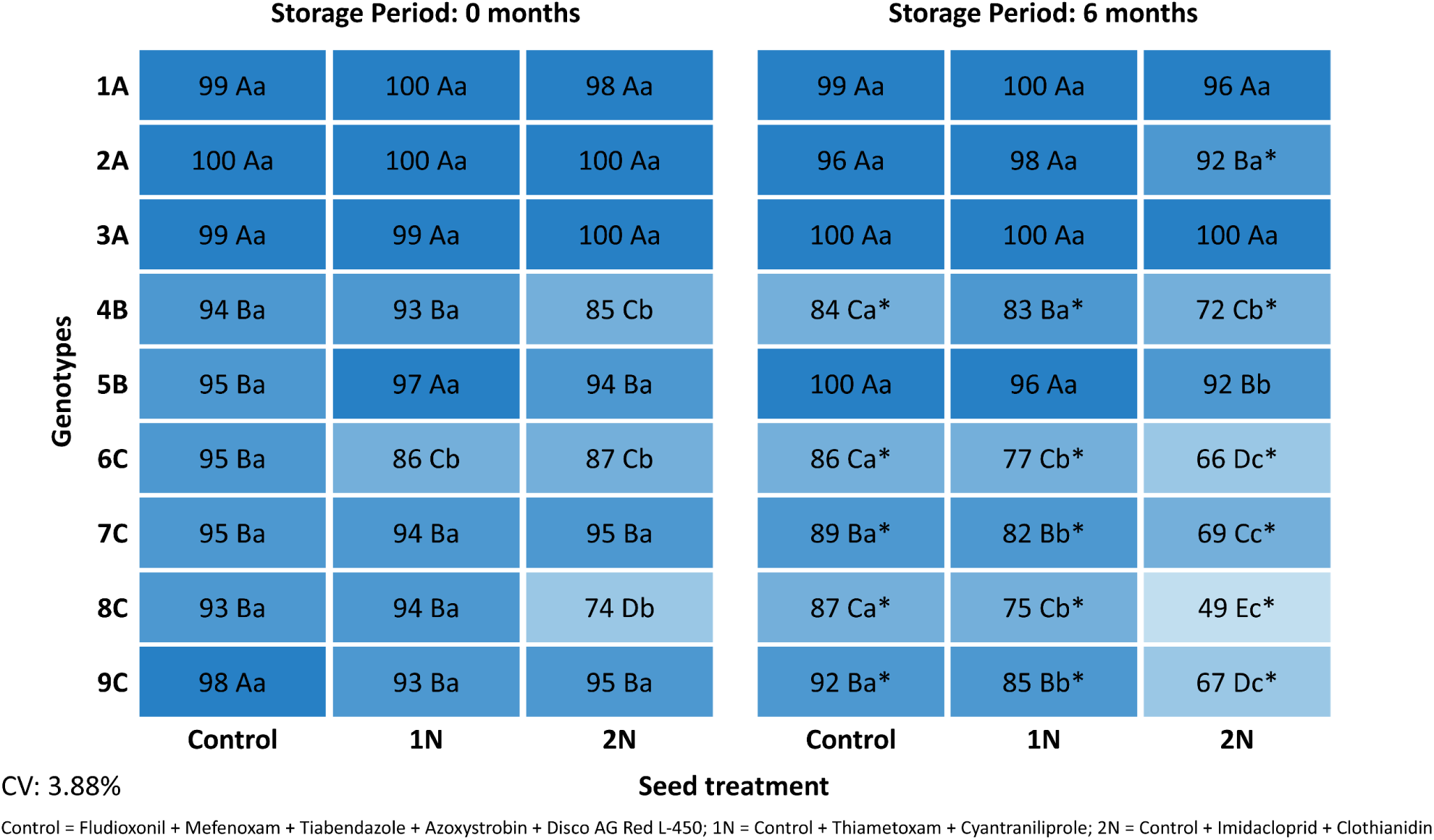
Percentage of emerged seedlings of maize in the cold test (7 days) as a function of genotype, seed treatment, and storage period (0 and 6 months at 25 °C). Statistical comparisons and notation as described in Figure 1.

### Phytotoxicity (Pi) Analysis

The phytotoxic effect of neonicotinoid insecticide treatments was quantified through the Pi. As observed for the physiological quality variables, a significant triple interaction among genotype, ST, and storage was detected in all tests.

At the beginning of storage, the RP test (Figure 5A) already revealed the sensitivity of certain genotypes: under 2N, 8C recorded a phytotoxicity index 9 pp higher than 1A. In the RP+V test (Figure 5B), this difference was even more pronounced, with female C genotypes and 4B recording indices at least 13 pp higher than female A genotypes in the same scenario. Under 1N, no difference among genotypes was observed in the RP+V test.

**Figure 5.**
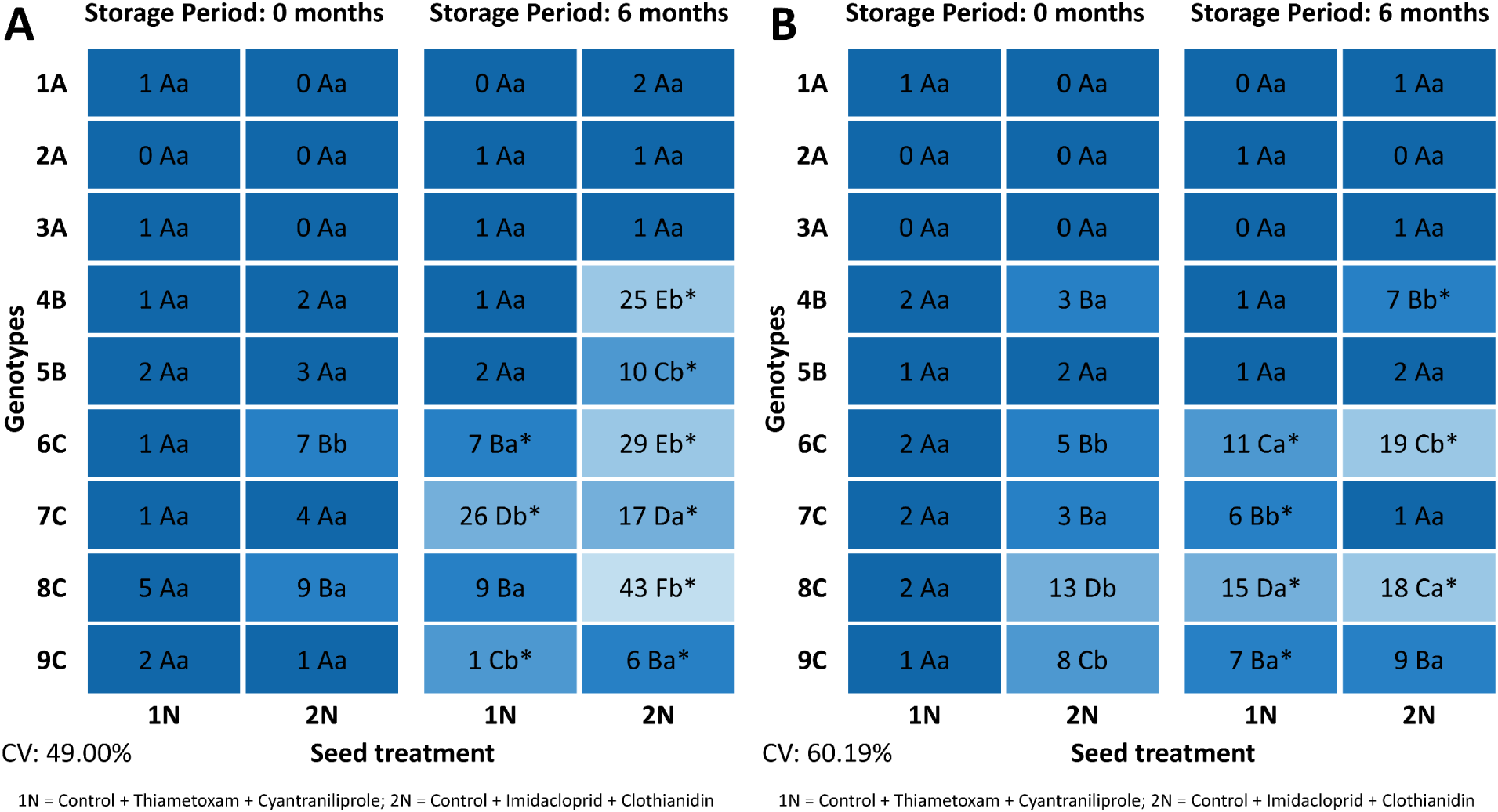
Phytotoxicity index (%) of maize seeds and seedlings in the (A) rolled paper germination test (7 days) and (B) rolled paper plus vermiculite germination test (7 days), as a function of genotype, seed treatment (1N and 2N), and storage period (0 and 6 months at 25 °C). The phytotoxicity index was calculated as the relative reduction in performance of insecticide treatments relative to the control for the same genotype and storage period. Statistical comparisons and notation as described in Figure 1.

After six months of storage, the phytotoxic effect was markedly amplified, especially in the RP test. Increases of 25 pp were observed for 7C (1N) and 34 pp for 8C (2N). At the end of the period, the maximum difference between the most sensitive and the most tolerant genotype was recorded under 2N, with 8C presenting an index 42 pp higher than 3A (Figure 5A). In the RP+V test, this difference was also observed but with lower intensity: while female A genotypes maintained phytotoxicity indices below 2%, 8C reached 18% (Figure 5B).

In the AA test (Figure 6A), the high sensitivity of female C genotypes was again highlighted. At the beginning of storage, 7C reached a phytotoxicity index 41 pp higher than 1A under 2N. After six months, phytotoxic damage increased by 80 pp for 6C and 75 pp for 9C under the same treatment. During this period, the sensitivity of female B genotypes also became evident, with 4B and 5B recording indices 35 and 51 pp higher than 3A under 2N, respectively.

**Figure 6.**
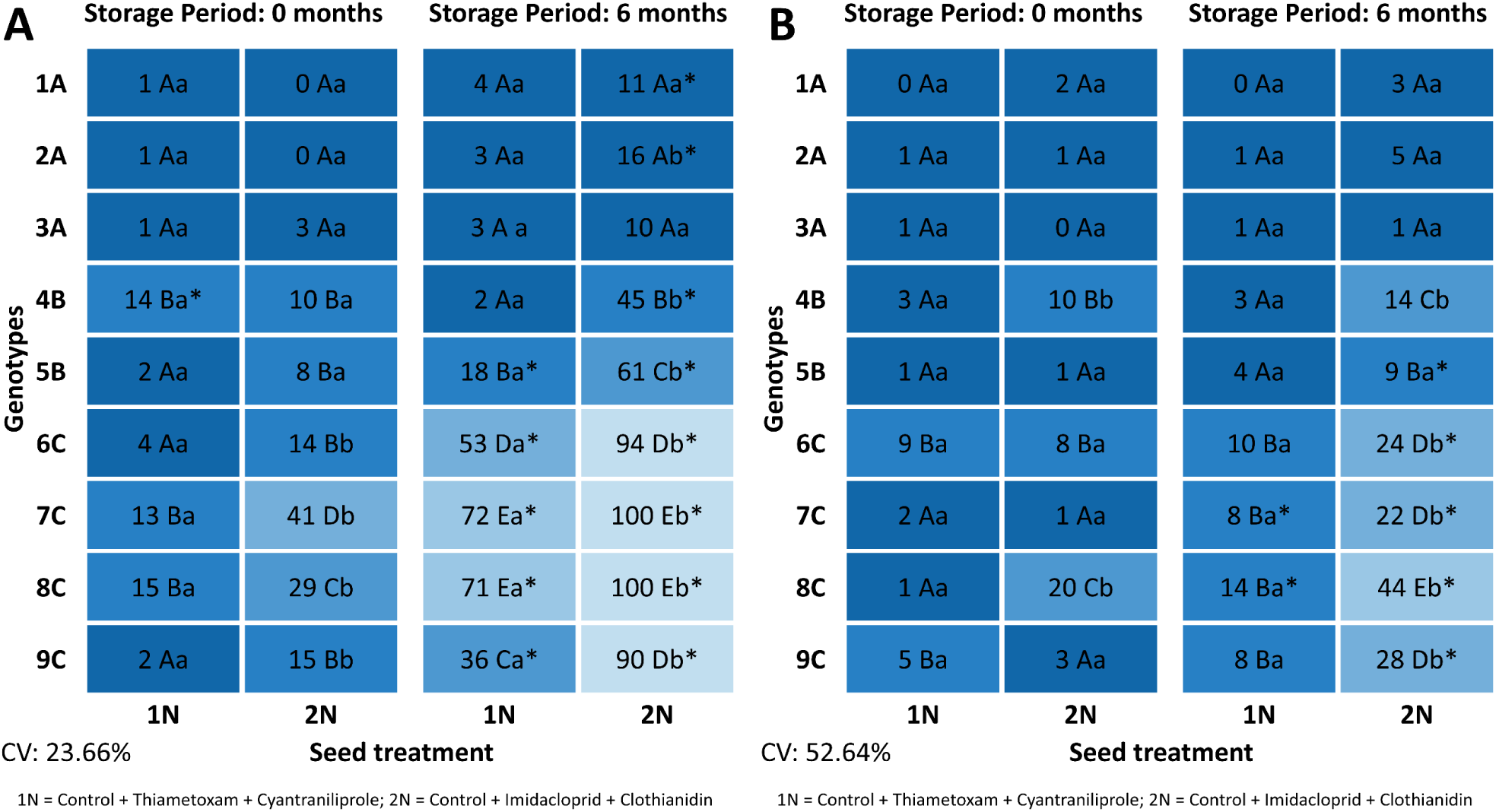
Phytotoxicity index (%) of maize seeds and seedlings in the (A) accelerated aging test (4 days) and (B) cold test (7 days), as a function of genotype, seed treatment (1N and 2N), and storage period (0 and 6 months at 25 °C). The phytotoxicity index was calculated as the relative reduction in performance of insecticide treatments relative to the control for the same genotype and storage period. Statistical comparisons and notation as described in Figure 1.

In the CT test (Figure 6B), the high sensitivity of 8C was confirmed, as it recorded the highest indices in nearly all scenarios. At the beginning of storage, its phytotoxicity index exceeded that of 3A by 20 pp under 2N, a difference that increased to 43 pp after six months. Storage also amplified phytotoxicity for other genotypes, such as 6C and 4B, with increases of 18 pp in the index. Treatment 2N was the most damaging after storage, raising the phytotoxicity of 9C to 28% compared to only 8% under 1N.

### Biochemical Analyses

To evaluate antioxidant enzyme activity, the most contrasting hybrids for phytotoxicity tolerance (3A, tolerant; 8C, sensitive) were selected. As with previous variables, a significant triple interaction among genotype, ST, and storage was observed for all analyses.

SOD activity revealed distinct patterns between genotypes (Figure 7A). At the beginning of storage, 2N reduced SOD activity in 3A by 20% relative to the control, while 8C maintained stable levels. After six months, storage induced an expressive increase in SOD activity under the control treatment, exceeding 60% for both genotypes. However, 2N maintained a suppressive effect on the enzyme during this period in both genotypes relative to the control.

**Figure 7.**
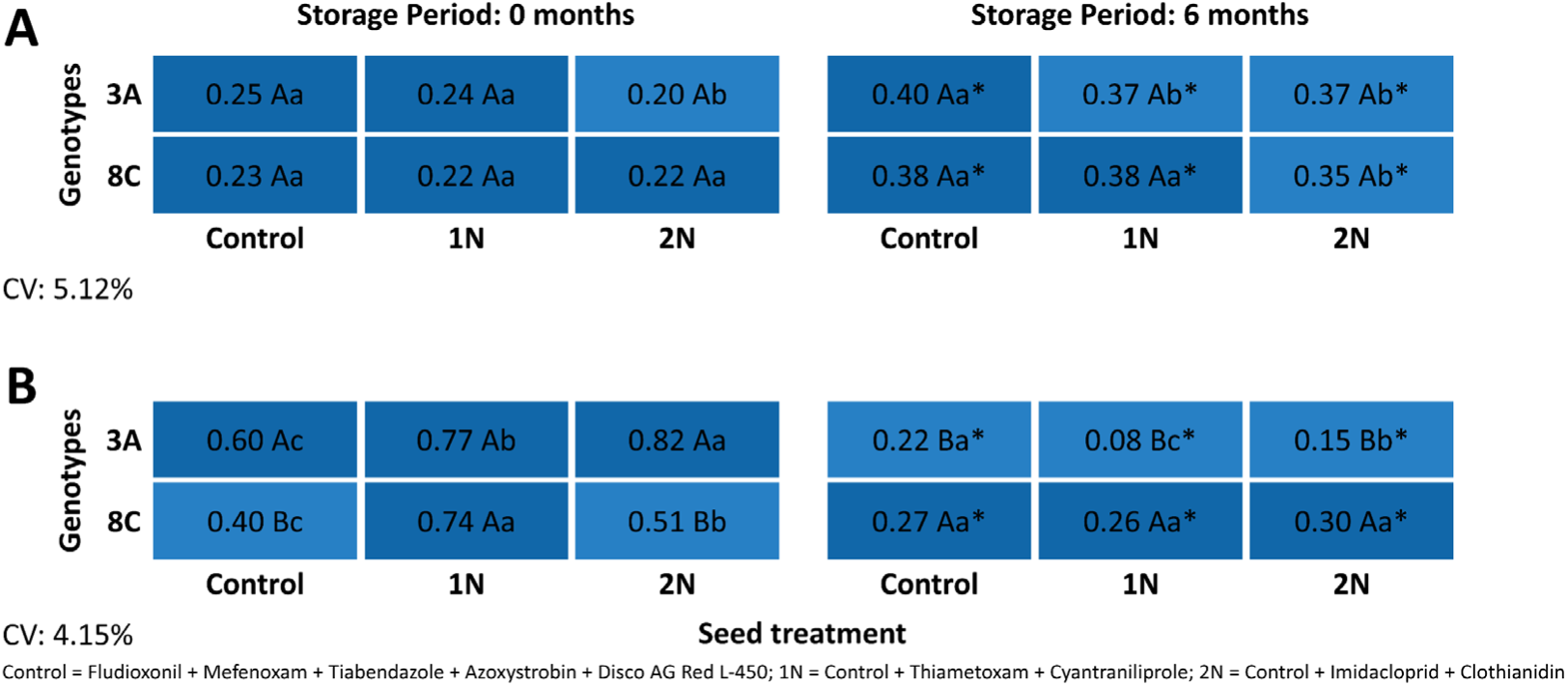
Enzymatic activity in maize seeds of the contrasting genotypes 3A (tolerant) and 8C (susceptible): (A) superoxide dismutase (SOD; U SOD mg⁻¹ fresh mass) and (B) catalase (CAT; µmol H₂O₂ min⁻¹ mg⁻¹ fresh mass), as a function of seed treatment (Control, 1N, and 2N) and storage period (0 and 6 months at 25 °C). Statistical comparisons and notation as described in Figure 1.

For CAT, a strong interaction among factors was verified, with behavioral inversion after storage (Figure 7B). Initially, 3A showed an active defensive response, with CAT activity more than 30% higher under 1N and 2N than in the control, surpassing 8C, which showed no response to 2N. After six months, a sharp decline in CAT activity was observed in 3A. In contrast, 8C, even with a reduction after storage, maintained higher and more stable CAT levels, becoming superior to 3A in CAT activity.

APX activity indicated a complex interaction between genotypes and stress intensity (Figure 8A). At the beginning of storage, 8C recorded higher activity under 1N (4.57 µmol ASA min⁻¹ mg⁻¹), while under the more severe stress of 2N, genotype 3A reached higher values than 8C. After six months, the combined effect of storage and 2N caused a drastic decline in APX activity in both genotypes, except in the control of 8C. However, the superiority of 3A was maintained, while 8C activity was reduced to residual levels (0.16 µmol ASA min⁻¹ mg⁻¹ under 2N).

**Figure 8.**
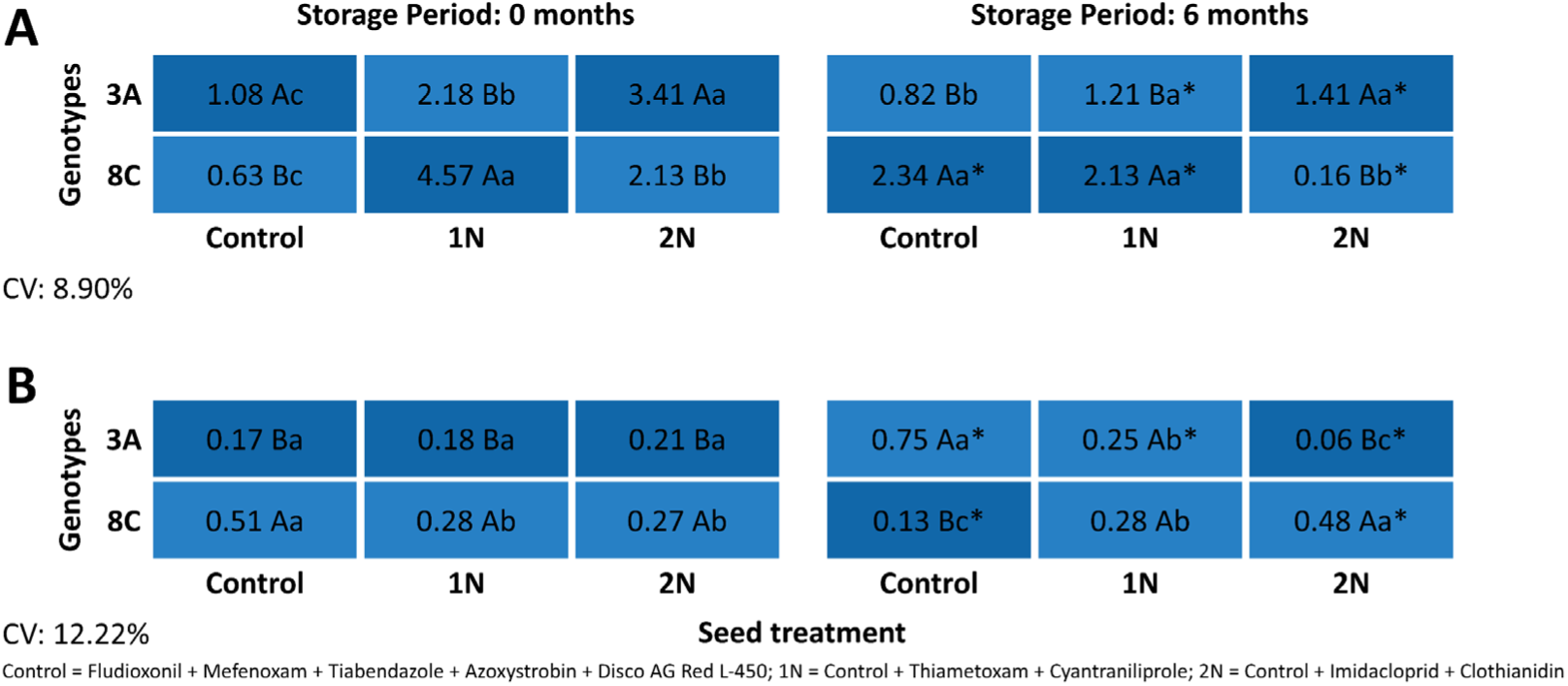
Enzymatic activity and hydrogen peroxide content in maize seeds of the contrasting genotypes 3A (tolerant) and 8C (susceptible): (A) ascorbate peroxidase (APX; µmol ASA min⁻¹ mg⁻¹ fresh mass) and (B) hydrogen peroxide (H₂O₂; mmol H₂O₂ mg⁻¹ fresh mass), as a function of seed treatment (Control, 1N, and 2N) and storage period (0 and 6 months at 25 °C). Statistical comparisons and notation as described in Figure 1.

H₂O₂ content also revealed contrasting patterns (Figure 8B). Initially, 8C accumulated more H₂O₂ in the control, indicating a higher basal oxidative stress, and was superior to 3A across all treatments. After six months, 3A reached the highest H₂O₂ accumulation in the control and the lowest under 2N, while 8C recorded the peak under the combination of storage and 2N.

### Genotype Classification Methodology Validation

The STTI synthesized genotype responses to chemical and storage stress (Figure S1, available in the Supporting Information). Female A genotypes (1A to 3A) showed high tolerance across all scenarios, while female C genotypes were more sensitive, especially 8C. Under 1N, STTI for 8C was reduced by 25.5% after six months, with greater sensitivity in the AA and CT tests (Figure S1A). Treatment 2N substantially intensified this loss, leading to a 45.7% reduction in STTI for 8C, including total collapse in AA (STTI = 0) (Figure S1B). In contrast, genotypes 1A to 3A maintained high tolerance, with STTI above 0.94 even in the most adverse scenario.

Multivariate analysis, combining STTI and Pi for each genotype across both storage periods, enabled separation into three distinct groups through hierarchical clustering (Figure 9). Classification became clearer after six months: Group 1 was formed by 1A, 2A, and 3A, which remained strongly clustered, indicating high stability. Group 3 included female C genotypes after six months, which shifted to extremely negative values on the first principal component (PC1) axis. These same genotypes, at the beginning of storage, occupied an intermediate position, indicating a progressive accentuation of sensitivity over time. PC1 explained 90.1% of the classification, and statistical distinction among groups was confirmed by MANOVA (Wilks’ Lambda = 0.010; F = 6.90; p < 0.001).

**Figure 9.**
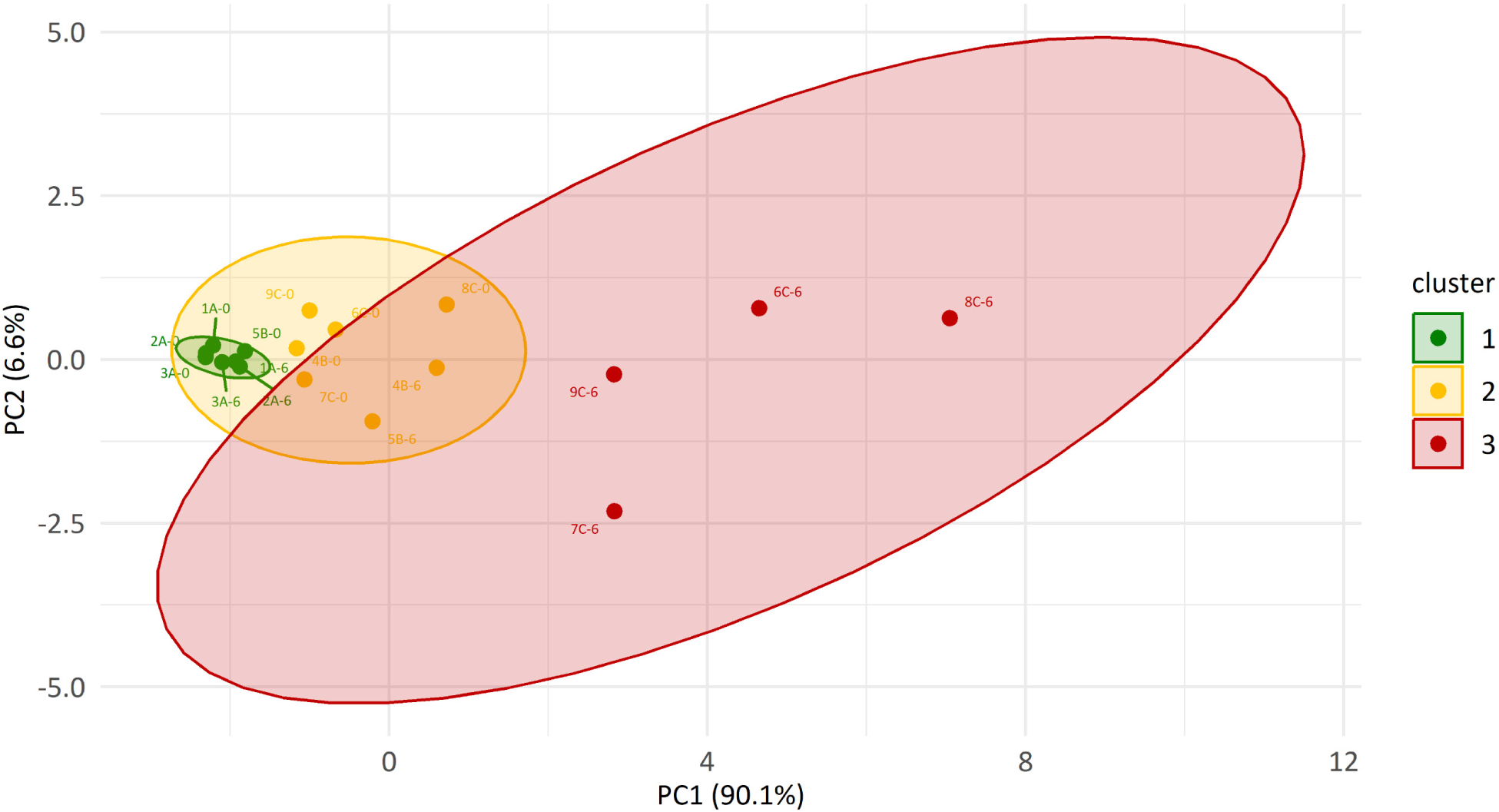
Principal component analysis (PCA) and hierarchical clustering (Ward’s method, Euclidean distance) of nine maize hybrid genotypes based on the Seed Treatment Tolerance Index (STTI) and phytotoxicity index (Pi) calculated across four physiological tests (RP, RPV, AA, and CT) at 0 and 6 months of storage at 25 °C. Symbols represent genotypes at 0 months (open) and 6 months (filled) of storage; ellipses indicate the three tolerance groups identified by cluster analysis and confirmed by MANOVA (Wilks’ Lambda = 0.010; F = 6.90; p < 0.001). PC1 explained 90.1% of total variance.

ANOVAs indicated significant differences (p < 0.001) among groups for all indices evaluated (Figure S2, available in the Supporting Information). Group 1 consistently recorded the lowest Pi values and the highest STTI close to 1, while Group 3 exhibited the opposite pattern. Differentiation among groups was maximized in the AA and CT vigor tests, where the Pi difference between Groups 1 and 3 reached 60 pp and the STTI of Group 3 was 58% lower than that of Group 1. Although both indices discriminated the groups, STTI revealed more consistent separation across the different physiological tests.

The final classification, based on the general STTI after six months, showed clear performance stratification among hybrids (Figure 10A). Genotypes 1A, 2A, and 3A composed the Tolerant group (STTI > 0.95). The Moderately Susceptible group included 4B and 5B (STTI between 0.80 and 0.89). The Susceptible group (6C, 7C, 8C, and 9C) showed greater internal variation, with STTI values between 0.73 and 0.57.

**Figure 10.**
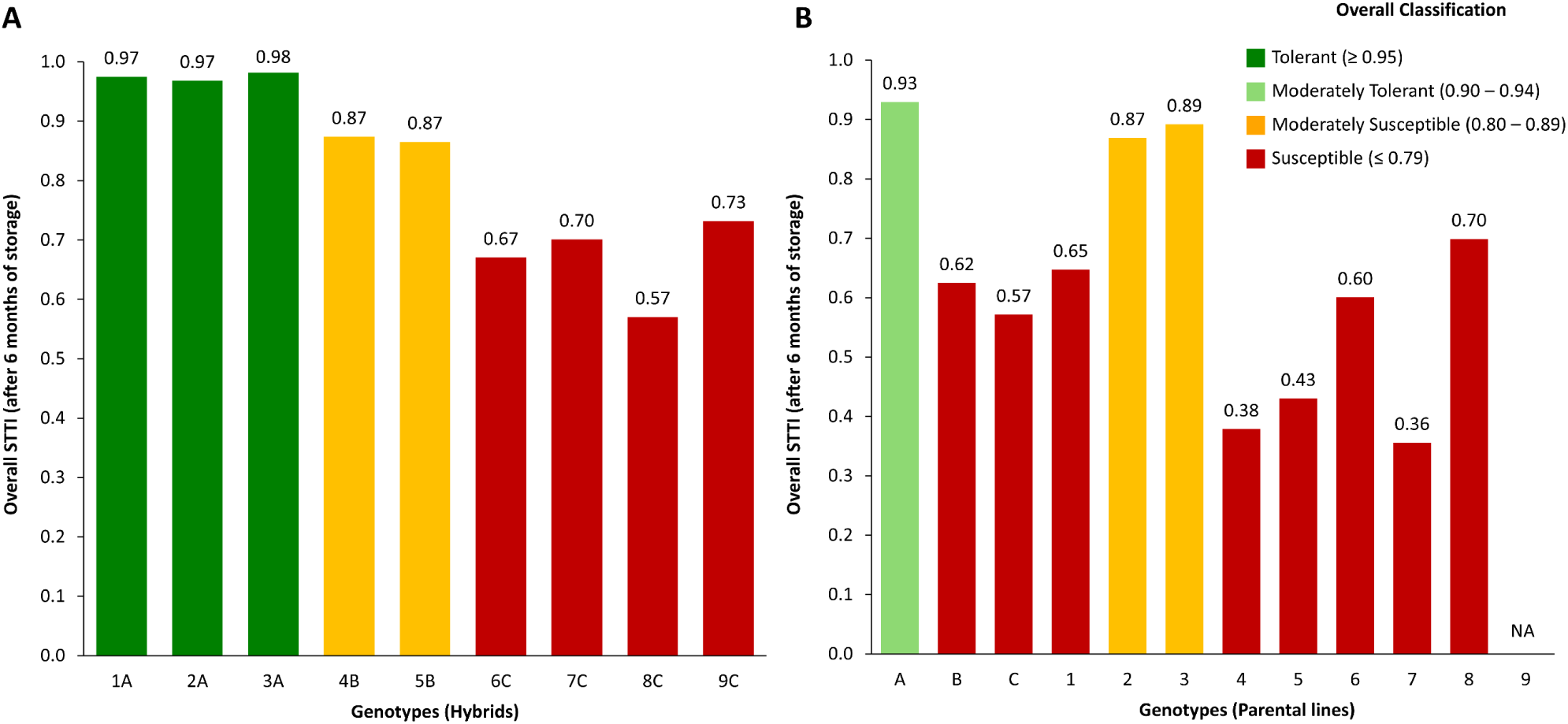
Classification of maize genotypes according to the general Seed Treatment Tolerance Index (STTI) after 6 months of storage at 25 °C: (A) commercial hybrids and (B) parental lines. The STTI was calculated as the mean ratio of physiological performance under insecticide treatments (1N and 2N) relative to the control across four tests (RP, RPV, AA, and CT). Horizontal dashed lines delimit tolerance groups: Tolerant (STTI > 0.95), Moderately Susceptible (0.80 ≤ STTI ≤ 0.89), and Susceptible (STTI < 0.80). Error bars represent standard deviation.

Parental lines revealed greater heterogeneity and generally lower tolerance than the hybrids (Figure 10B). The most tolerant (female A lines) reached STTI close to 0.90, while the most susceptible (lines 4, 5, and 7) recorded values below 0.45. Most parental lines were positioned in the susceptible range, with indices lower than those of hybrids in the same susceptible group.

## DISCUSSION

The results obtained in this study confirm that maize seed treatment with neonicotinoid insecticides compromises physiological quality and that storage at 25 °C for six months accentuates this effect in susceptible genotypes. Among the seed treatments evaluated, 2N was the most phytotoxic because it contains two neonicotinoids, amplifying genotypic differences over storage. Hybrids derived from female A maintained high performance with minimal physiological quality losses, while those from female C suffered the most, especially in the AA and CT vigor tests (Krzyzanowski, 2020), which proved to be the most sensitive tests for differentiating STTI. These findings are reinforced by the multivariate analysis, which robustly separated genotypes into tolerance groups with a consistent pattern of alignment among hybrid performance.

Understanding these results requires considering the mechanisms by which neonicotinoids exert phytotoxicity on seeds. Although widely studied for their efficacy in controlling early-season pests (Neves et al., 2022; Ismail, 2024), their physiological effects on seeds and seedlings remain insufficiently explored (Reis et al., 2026a). Phytotoxicity from seed treatment manifests through several interconnected mechanisms, with oxidative stress induction as the primary damage pathway (Forti et al., 2024). This oxidative imbalance leads to seed deterioration: in freshly treated seeds, constitutive antioxidants and repair mechanisms may partially contain such damage, but residual toxicity tends to persist during storage, favoring the progression of deterioration (Oliveira et al., 2020). Reis et al. (2026a) proposed a three-tier classification of phytotoxicity severity based on cellular damage progression, ranging from limited damage (recoverable vigor reduction) to extreme damage (seed death), with neonicotinoids capable of causing all three levels depending on slurry composition, storage duration, and genotypic tolerance.

This interaction is particularly relevant because the physiological deterioration process, characterized by lipid peroxidation, combined with neonicotinoid effects, can compromise membrane integrity, alter cell permeability, and impair mitochondrial function, affecting the energy required for germination (Liu et al., 2021). Transcriptomic studies in maize seedlings exposed to imidacloprid revealed increased lipid peroxidation, alterations in genes related to antioxidants and detoxification (glutathione S-transferases, ABC transporters), and reduced growth associated with oxidative damage (Zhang et al., 2022). When combined with chemical stress, a synergistic condition forms: the seed begins storage with initial damage and, over time, the declining antioxidant capacity potentiates the impact of the neonicotinoid insecticide (Oliveira et al., 2020). This interaction corroborates the observations of Reis et al. (2026b), Medeiros et al. (2026) and Moraes et al. (2022), who also reported reductions in maize seed quality treated with neonicotinoids after 180 days of storage. In this study, such deterioration was even more precocious: susceptible genotypes already reflected phytotoxicity when freshly treated, and after six months, damage was severely amplified, with reductions of up to 48 pp in germination and 90 pp in vigor (AA) under 2N. Tolerant hybrids, by contrast, showed none of these behaviors, confirming that the genetic constitution is the determining factor for a maize seed’s capacity to withstand the phytotoxic effects of insecticide treatment over storage.

The biochemical evaluation revealed that the tolerance observed in physiological tests is strongly associated with the capacity to control H₂O₂ accumulation. In tolerant genotypes such as 3A, even under 2N and storage, H₂O₂ remained at low levels, sustained by relatively stable APX and CAT activity. Sofo et al. (2015) observed that tolerant cultivars of *Jatropha curcas* under drought maintain stable CAT and APX activities, avoiding H₂O₂ accumulation, as also observed in genotype 3A. These antioxidant enzymes function in H₂O₂ detoxification (Anjum et al., 2016). Although CAT activity declined over time, APX played a compensatory role, ensuring H₂O₂ detoxification and retarding deterioration. According to Anjum et al. (2016) CAT and APX act complementarily: CAT responds to high H₂O₂ peaks, while APX ensures continuous elimination at low to moderate concentrations. When there is a compensatory increase in APX and a decline in CAT, this is indicative of greater tolerance to a given stress (Anjum et al., 2017).

In susceptible genotypes, expressive H₂O₂ accumulation was accompanied by marked reductions in CAT and APX, resulting in low efficiency of the main detoxification routes. This condition likely led to loss of cellular integrity and accelerated deterioration (Hasanuzzaman et al., 2020). Forti et al. (2024) measured H₂O₂ increases of 67% above the control and a CAT activity reduction of 250% relative to the control in maize seedlings exposed to thiamethoxam, demonstrating the phytotoxic potential of this neonicotinoid. It is important to note that SOD activity showed little variation between genotypes or treatments, indicating that O₂⁻ conversion to H₂O₂ was not the limiting factor, as also observed by Saed-Moucheshi et al. (2021) in triticale. Therefore, the critical point resides in the efficient elimination of H₂O₂ before it causes irreversible oxidative damage, a central task of CAT and APX (Anjum et al., 2016).

This critical dependence on H₂O₂ elimination explains why the greatest genotypic differences were revealed in the AA and CT vigor tests. By adding environmental stress to the pre-existing chemical stress, these tests expose the deficiencies in the antioxidant defense system of each hybrid. The AA test intensifies ROS generation by subjecting seeds to high temperature and humidity, overloading the detoxification capacity (Marcos-Filho, 2020). The CT test directly challenges membrane integrity and respiratory capacity under low temperature (Cicero and Vieira, 2020). In this scenario, susceptible hybrids with elevated residual H₂O₂ levels likely failed to maintain ionic gradients and the energy required for emergence under cold stress. This explains the increase in phytotoxicity observed even in hybrids considered tolerant under the ideal conditions of the RP and RP+V germination tests, where they maintained their performance regardless of treatment or storage, as those tests impose only chemical stress without the additional environmental challenge.

The integration of results through the multivariate approach reinforced the interpretation of genotypic differences. The STTI synthesized multiple physiological parameters into a single index, while hierarchical clustering and PCA revealed consistent response patterns, validated by MANOVA. The multivariate analysis made the distinction among hybrids regarding phytotoxicity tolerance even clearer, particularly when evaluated at 6 months of storage. This distinction is manifested, in practice, by the significant increase in phytotoxicity over storage observed in female B and C hybrids, indicating an intrinsic inability of these genotypes to mitigate cumulative cellular damage, which evolves into intensified physiological losses (Reis et al., 2026a). Our findings are consistent with those of Deuner et al. (2014) who also reported a 16 pp reduction in germination of maize seeds treated with thiamethoxam, albeit over 12 months of storage. The STTI is therefore a decision-support tool with direct practical implications for the seed chain. For breeding programs, it enables efficient screening and selection of genotypes with greater tolerance to seed treatment and storage. For the seed industry, it provides objective criteria to optimize logistics by defining the ideal seed treatment window and the best decision-making strategy for choosing stored lots based on the STTI classification of each hybrid, as illustrated by the classification proposed in Figure 10.

The consistent superiority of hybrids derived from female A points to a probable maternal component in the inheritance of tolerance, given that this line is classified as moderately tolerant by the proposed classification. Although confirmation of the maternal inheritance hypothesis requires future studies with reciprocal crosses, the mechanisms may involve a greater initial supply of antioxidants and reserves from the maternal plant, or inherited structural characteristics such as a more robust pericarp that limits insecticide absorption at the time of ST (Gong et al., 2024; Brunel-Muguet et al., 2025; Reis et al., 2026b). Additionally, epigenetic mechanisms, such as DNA methylation patterns or regulation by small RNAs, which may modulate stress response gene expression in a transgenerational manner, cannot be ruled out (Chow and Mosher, 2023). Several studies point to genetic constitution as a determining factor in high physiological quality and stress response (Marques et al., 2019; Costa et al., 2021). Reinforcing this premise, Melnik et al. (2023) found, in a study with reciprocal crosses, that the maternal effect is a key factor in maize seed quality, with certain parental lines conferring superior quality to hybrids only when used as the female progenitor, and that this performance difference was further amplified under stress conditions such as the cold test.

Despite the relevance of the findings, important limitations must be recognized, which open avenues for future research. The biochemical analysis was focused on contrasting genotypes, and extrapolation to a broader genotype spectrum should be made with caution. Quantification of insecticide residues in seed embryonic tissues (Alford and Krupke, 2017) and classical damage biomarkers such as malondialdehyde (Shahid et al., 2021) would strengthen the causal connection between internal chemical dose and the magnitude of oxidative injury. For more applied research, the expansion of the genetic panel, the inclusion of different active ingredient combinations, and the evaluation of seed performance under other storage conditions are recommended. Large-scale genotype screening can be accelerated by the use of rapid, non-destructive phenotyping techniques, such as multispectral image analysis (ElMasry et al., 2019). Parallel studies focused on the maternal effect and the genetic bases of tolerance are essential to direct breeding programs more effectively (Melnik et al., 2023). From an operational perspective, STTI application can be adapted to other industrially treated crops, broadening the practical impact of this methodology.

Tolerance of maize seeds to treatment and storage is a characteristic dependent on genetic constitution. Susceptibility manifests early in sensitive genotypes but is amplified by storage. The biochemical mechanism of tolerance resides in the differential capacity to regulate H₂O₂, whereby tolerant genotypes maintain homeostasis through efficient catalase and ascorbate peroxidase activity. The STTI proved to be a high-discriminatory tool, recommended for evaluation at 6 months post-storage. This approach, together with the proposed classification, represents an advance for large-scale screening in genetic breeding programs and offers an objective criterion for optimizing seed treatment logistics based on the specific tolerance of each genotype.

## ABBREVIATIONS USED

AA: Accelerated Aging test
ANOVA: Analysis of Variance
APX: Ascorbate Peroxidase
CAT: Catalase
CT: Cold Test
DNA: Deoxyribonucleic Acid
H₂O₂: Hydrogen Peroxide
MANOVA: Multivariate Analysis of Variance
PC1: First Principal Component
PCA: Principal Component Analysis
ROS: Reactive Oxygen Species
RP: Rolled Paper germination test
RP+V: Rolled Paper plus Vermiculite
SOD: Superoxide Dismutase
STTI: Seed Treatment Tolerance Index
ST: Seed Treatment

## AUTHOR INFORMATION

## Author Contributions

V.U.V. Reis: Conceptualization, data curation, formal analysis, investigation, methodology, visualization, writing (original draft), writing (review and editing); G.I.S. Tavares: Investigation, writing (review and editing); D.C. Maciel: Investigation, writing (review and editing); J.P. Januario: Investigation (biochemical analyses); M.S.R. Pereira: Investigation, writing (review and editing); R.M.O. Pires: Resources, writing (review and editing); E.R. Carvalho: Conceptualization, funding acquisition, project administration, supervision, writing (review and editing). All authors have given approval to the final version of the manuscript.

## Funding Sources

Conselho Nacional de Desenvolvimento Científico e Tecnológico, Grant No. 307200/2025-6; Fundação de Amparo à Pesquisa do Estado de Minas Gerais; Coordenação de Aperfeiçoamento de Pessoal de Nível Superior, Finance code 001.

## Conflict of interests

The authors declare no competing financial interest.

## ASSOCIATED CONTENT

The following files are available free of charge.

Supporting_Information.pdf: Detailed experimental protocols for physiological and biochemical analyses, including germination and vigor tests, antioxidant enzyme assays (SOD, CAT, and APX), hydrogen peroxide quantification, phytotoxicity index (Pi) calculation, Seed Treatment Tolerance Index (STTI) formula, and statistical procedures; Tables S1 and S2 with genotype crosses and seed treatment product descriptions; Figure S1: radar plots of the STTI for all nine maize hybrid genotypes under 1N and 2N treatments across storage periods (0 and 6 months); Figure S2: boxplot comparison of Pi and STTI among tolerance groups with ANOVA results confirming significant differences for all evaluated physiological tests.

## ACKNOWLEDGMENTS

The authors extend sincere gratitude to the Conselho Nacional de Desenvolvimento Científico e Tecnológico (CNPq), the Fundação de Amparo à Pesquisa do Estado de Minas Gerais (FAPEMIG), and the Coordenação de Aperfeiçoamento de Pessoal de Nível Superior (CAPES) for their generous financial support, including scholarships and a research productivity grant. Thanks are also extended to Syngenta and Seedcare Institute (Brazil) for the partnership.

## Notes

### Competing Interest Statement

The authors have declared no competing interest.

