## Supporting Information for "Phenotyping maize seed tolerance to storage after seed treatment using a Seed Treatment Tolerance Index"

### Potential methodology for phenotyping maize seed tolerance to storage after seed treatment

This document contains detailed experimental protocols and additional figures referenced in the main manuscript.

#### SUPPLEMENTARY MATERIALS AND METHODS

##### *S1. Genotypes*

Nine commercial maize (*Zea mays* L.) hybrids and their respective parental lines from the 2022/2023 harvest season were used (Table S1). To preserve commercial confidentiality, genotypes were coded. After harvest, ears were dried in a dryer to 12% moisture content, seeds were processed, and subsequently delivered to the seed analysis laboratory.



**Table S1.** Crosses of the genotypes used.

| Hybrid | Male parent | Female parent |
|---|---|---|
| 1A | 1 | |
| 2A | 2 | A |
| 3A | 3 | |
| 4B | 4 | B |
| 5B | 5 | |
| 6C | 6 | |
| 7C | 7 | C |
| 8C | 8 | |
| 9C | 9 | |

*S2. Seed Treatments and Storage Conditions*

Seeds were treated in 2-kg batches using an Arktos Laboratório L2K-BM industrial-batch simulator (Momesso, Matão, Brazil), with the frequency inverter calibrated at 15 Hz to simulate industrial batch-type treatment. The complete application and homogenization process for each batch lasted 20 seconds.

Seeds of each genotype were subjected to three treatments (Table S2):

- **Control:** treatment with fungicide and polymer.
- **1N:** control treatment supplemented with one neonicotinoid insecticide and one diamide insecticide.
- **2N:** control treatment supplemented with a mixture of two neonicotinoid insecticides.



**Table S2.** Slurry composition of seed treatments applied to the genotypes.

| Seed treatment | Active ingredient (AI) | Class | AI dose† | Commercial product dose† |
|---|---|---|---|---|
| **Control** | Azoxystrobin | Fungicide | 0.04 g | 25 mL |
| | Thiabendazole | Fungicide | 7.50 g | |
| | Fludioxonil | Fungicide | 0.94 g | |
| | Metalaxyl-M | Fungicide | 0.75 g | |
| | - | Polymer* | - | 40 mL |
| **1N** | **Control components +** | | | |
| | Cyantraniliprole | Diamide insecticide | 4.20 g | 70 mL |
| | Thiamethoxam | Neonicotinoid insecticide | 3.60 g | 60 mL |
| **2N** | **Control components +** | | | |
| | Imidacloprid | Neonicotinoid insecticide | 3.60 g | 60 mL |
| | Clothianidin | Neonicotinoid insecticide | 4.20 g | 70 mL |

†Dosage in g or mL per 60,000 seeds. *Polymer used: Disco AG Red L-450® (density 1.05–1.15 g $mL^{-1}$, viscosity 300–1000 cPs at 25 °C, pH 6–8).

Following treatment, 1 kg of seeds from each batch was designated for evaluations at time zero, while the remainder was stored for six months in a Biochemical Oxygen Demand (B.O.D.) incubator, model TH.7000 Touch (Thoth Equipamentos, Piracicaba, Brazil), at 25 ± 2 °C to simulate uncontrolled storage conditions in tropical/subtropical environments. Evaluations were conducted at two time points: immediately after treatment (time zero) and at the end of the six-month storage period.



*S3. Experimental Design*

The study was conducted in a completely randomized design. Three distinct factorial arrangements were used according to the nature of each evaluation:

*Physiological evaluations*: A 9 (hybrids) × 3 (treatments: Control, 1N, 2N) × 2 (storage periods: 0 and 6 months) factorial arrangement with four replications. Each experimental unit consisted of 50 seeds.

*Phytotoxicity*: For the calculation of the phytotoxicity index, comparing insecticide treatments against the control, a 9 (hybrids) × 2 (treatments: 1N, 2N) × 2 (storage periods: 0 and 6 months) factorial arrangement with four replications was used. Each experimental unit consisted of 50 seeds.

*Biochemical analyses*: For antioxidant enzyme evaluation, a 2 (contrasting hybrids based on physiological evaluations) × 3 (treatments: Control, 1N, 2N) × 2 (storage periods: 0 and 6 months) factorial arrangement with three replications was used. Each experimental unit consisted of 200 mg of macerated seeds.

*S4. Physiological quality evaluation*

Physiological quality evaluations were conducted following the protocols described below:

*Rolled paper germination test* (RP): Seeds were placed on a paper towel sheet moistened with distilled water at 2.5 times the paper mass. Seeds were then covered with another germination paper sheet, and rolls were assembled. Rolls were maintained in a Mangelsdorf-type germinator, model 4001 (Biomatic, Porto Alegre, Brazil), at 25 ± 2 °C. Evaluation was performed at seven days by recording the percentage of normal seedlings (Brasil, 2025).

*Rolled paper plus vermiculite germination test* (RP+V): After placing seeds on a moistened germination paper sheet (3× paper mass), 100 mL of moist vermiculite (1:1 v/v ratio) was added



and distributed uniformly over the seeds. Seeds were then covered with another germination paper sheet and rolls assembled, maintained under the same conditions as the RP test. Evaluation was performed at seven days by recording the percentage of normal seedlings (Rocha et al., 2023).

*Accelerated aging test* (AA): Plastic boxes with suspended aluminum screens were used. A uniform layer of seeds was distributed over the screens, with 40 mL of water at the bottom of each box. Boxes were placed in a B.O.D. chamber, model EL102 G (Eletrolab, São Paulo, Brazil), at 43 ± 1 °C for 72 h. After this period, seeds were sown following the germination test protocol, with evaluation of normal seedlings at four days after sowing, expressed as a percentage (Marcos-Filho, 2020).

*Cold test* (CT): Seeds were sown in plastic trays (51 × 30 × 9 cm) containing a sand-and-soil substrate (2:1 v/v ratio), moistened to 60% of water retention capacity. Trays were maintained in a cold chamber at 10 ± 2 °C for seven days and then transferred to a plant growth chamber at 25 ± 2 °C with a 12 h light/dark photoperiod for an additional seven days. At the end, the percentage of emerged seedlings was determined (Cicero and Vieira, 2020).

*S5. Antioxidant Enzyme Analyses and Hydrogen Peroxide Quantifications*

Seed samples (10 g) were macerated in liquid nitrogen with the addition of insoluble polyvinylpolypyrrolidone (PVPP; 50 mg $g^{-1}$ fresh tissue) to minimize oxidation of phenolic compounds. The pulverized material was subdivided into microtubes (200 mg each) and stored in an ultra-freezer at −80 °C, model MDF-U52VAT (Sanyo Electric Co. Ltd., Moriguchi, Japan), until analysis. Readings were performed in triplicate using an Eon™ spectrophotometer (BioTek Instruments, Winooski, USA).

*S5.1. Antioxidant Enzymes*



*Crude enzymatic extract preparation*: Samples of 200 mg were homogenized in 1.5 mL of 100 mM potassium phosphate buffer (pH 7.8) containing 0.1 mM EDTA and 10 mM ascorbic acid. The mixture was centrifuged at 13,000 × g for 10 min at 4 °C in a refrigerated centrifuge, model SL-706 (Solab, Piracicaba, Brazil), and the supernatant was used for enzymatic activity determinations.

*SOD activity*: Determined according to Giannopolitis & Ries (1977), based on the enzyme's ability to inhibit the photoreduction of nitro-blue tetrazolium (NBT). The assay was performed in 96-well plates with a final volume of 200 µL per reaction, containing: 20 µL of enzymatic extract, 100 µL of 100 mM potassium phosphate buffer (pH 7.8; 50 mM final concentration), 40 µL of 70 mM methionine (14 mM final), 3 µL of 10 µM EDTA (0.1 µM final), 21 µL of distilled water, 15 µL of 1 mM NBT (75 µM final), and 2 µL of 0.2 mM riboflavin (2 µM final). NBT and riboflavin were added to the buffer immediately before the reaction, protected from light. The reaction was initiated by riboflavin addition and subsequent exposure to fluorescent light (30 W) for 7 min. Absorbance was recorded at 560 nm. One unit of SOD (U) was defined as the amount of enzyme required to inhibit 50% of NBT photoreduction under assay conditions. Results were expressed as U SOD $mg^{-1}$ fresh mass (FM).

*CAT activity*: Determined according to Azevedo et al., (1998) with adaptations from Havir & McHale (1987). The assay was conducted in half-area 96-well plates with a final volume of 180 µL per reaction, containing: 9 µL of enzymatic extract, 90 µL of 200 mM potassium phosphate buffer (pH 7.0; 100 mM final), and 72 µL of distilled water, pre-incubated in a water bath at 30 °C. Immediately before reading, 9 µL of 250 mM hydrogen peroxide ($H_2O_2$; 12.5 mM final) was added. The reaction was initiated by $H_2O_2$ addition, and decomposition was monitored at 240 nm for 3 min with readings every 15 s, using a molar extinction coefficient of 36 $mM^{-1}$ $cm^{-1}$.



Activity was expressed as µmol $H_2O_2$ decomposed per minute per mg fresh mass (µmol $H_2O_2$ $min^{-1}$ $mg^{-1}$ FM).

*APX activity*: Determined according to Nakano & Asada (1981), in half-area 96-well plates with a final volume of 180 µL per reaction, containing: 9 µL of enzymatic extract, 90 µL of 200 mM potassium phosphate buffer (pH 7.0; 100 mM final), 9 µL of 10 mM ascorbic acid (0.5 mM final), and 63 µL of distilled water, pre-incubated in a water bath at 30 °C. The reaction was initiated by addition of 9 µL of 2 mM $H_2O_2$ (0.1 mM final), and ascorbate oxidation was monitored at 290 nm for 3 min with readings every 15 s, using a molar extinction coefficient of 2.8 $mM^{-1}$ $cm^{-1}$. Activity was expressed as µmol of oxidized ascorbate (ASA) per minute per mg fresh mass (µmol ASA $min^{-1}$ $mg^{-1}$ FM).

*S5.2. Hydrogen Peroxide*

Quantification was performed according to Velikova et al., (2000) with adaptations as described below.

*Crude extract preparation*: Samples of 200 mg fresh mass were homogenized in 1.5 mL of 0.1% (w/v) trichloroacetic acid (TCA). The homogenate was centrifuged at 12,000 × g for 15 min at 4 °C, and the supernatant was collected for analysis.

*Standard curve*: Prepared from a 250 µM $H_2O_2$ stock solution, diluted to final concentrations of 0, 5, 15, 25, 35, and 45 µmol. For each point, 45 µL of 10 mM potassium phosphate buffer (pH 7.0), 90 µL of 1 M potassium iodide (KI), and the corresponding volume of $H_2O_2$ solution were mixed, with distilled water added to 45 µL total.

*$H_2O_2$ quantification*: For each sample, 45 µL of supernatant, 45 µL of 10 mM potassium phosphate buffer (pH 7.0; 2.5 mM final), and 90 µL of 1 M KI (0.5 M final) were mixed. Samples were kept in the absence of light, and absorbance was recorded at 390 nm. $H_2O_2$



concentration was calculated based on the standard curve and expressed as mmol $H_2O_2$ mg$^{-1}$ fresh mass.

*S6. Index Calculations*

Index calculations were performed using data from the physiological quality evaluations.

*Phytotoxicity index* (Pi): Calculated for each physiological variable relative to the control treatment (seeds without insecticide, same genotype and storage period). The variable expresses the percentage reduction of the value observed under insecticide treatments relative to the control mean, according to Equation S1 (Reis et al., 2026):

$$Pi\ (\%) = \frac{(\bar{C} - T)}{\bar{C}} \times 100 \qquad \text{(S1)}$$

where $\bar{C}$ is the mean of the analyzed variable for the control treatment in the same genotype and storage period, and $T$ is the value of one replicate of the analyzed variable for the insecticide treatment.

Positive values indicate a phytotoxic effect. Negative values, interpreted as absence of phytotoxicity, were adjusted to zero prior to statistical analyses. This approach allowed comparison of the relative impact of insecticide treatments among genotypes, minimizing the influence of intrinsic performance differences.

*Seed Treatment Tolerance Index* (STTI): Calculated for each combination of genotype, treatment, and storage period across all physiological variables evaluated. For each variable, the index was obtained as the ratio of the mean observed under the insecticide treatment (1N or 2N) to the corresponding control mean, according to Equation S2:

$$STTI = \frac{\bar{\bar{T}}}{\bar{\bar{C}}} \qquad \text{(S2)}$$



where $\overline{T}$ is the mean of the variable under insecticide treatment (1N or 2N) and $\overline{C}$ is the mean of the variable under the control treatment (same genotype and storage period).

STTI values were then calculated separately for each physiological variable and storage period. To integrate the overall genotype response, the arithmetic mean of STTI values obtained across all physiological variables and both treatments (1N and 2N) was used, yielding a single tolerance index per genotype.

STTI normalization reduces the influence of intrinsic differences in natural deterioration among genotypes, enabling relative comparisons of seed treatment impact and facilitating genotype classification.

*S7. Statistical Analysis*

Data analysis was conducted in R studio software in two main stages (R Core Team, 2024).

*S7.1. Evaluation of Experimental Effects*

To evaluate the main effects of the factors genotype, seed treatment, and storage period, as well as their interactions, individual analysis of variance (ANOVA) was performed for each variable from the physiological quality tests, phytotoxicity indices (Pi), and antioxidant enzyme activities. Treatment means with significant effects ($\alpha < 0.05$) in ANOVA were compared using the Scott–Knott test ($\alpha < 0.05$) with the ExpDes.pt package (Ferreira et al., 2021).

*S7.2. Development and Validation of the Classification Methodology*

Genotype classification for tolerance was based on sequential multivariate analysis, using Pi and STTI as input variables, with analyses performed in the MVar.pt package (Ossani and Cirillo, 2025).

The analytical sequence was as follows:

*Data preparation*: Pi and STTI values were organized into a matrix containing experimental unit identifiers (ID). To remove replicates, data were grouped by ID and the arithmetic mean of numerical variables was calculated for each ID.

*Clustering and visualization*: The aggregated matrix was standardized (z-score transformation) and submitted to hierarchical cluster analysis using Ward's method with Euclidean distance (Ward, 1963). Cluster visualization was performed through principal component analysis (PCA).

*Group validation*: Statistical distinction among clusters was verified by multivariate analysis of variance (MANOVA) using Wilks' Lambda criterion (Wilks, 1932), followed by univariate ANOVAs for each variable to identify those contributing most to group separation.

*Final classification*: Based on the results, the final tolerance classification of genotypes to storage after seed treatment was established from the mean of all STTI variables, for both insecticide treatments, at six months of storage. STTI was also calculated for parental lines, although only the general value is presented.



### SUPPORTING FIGURES

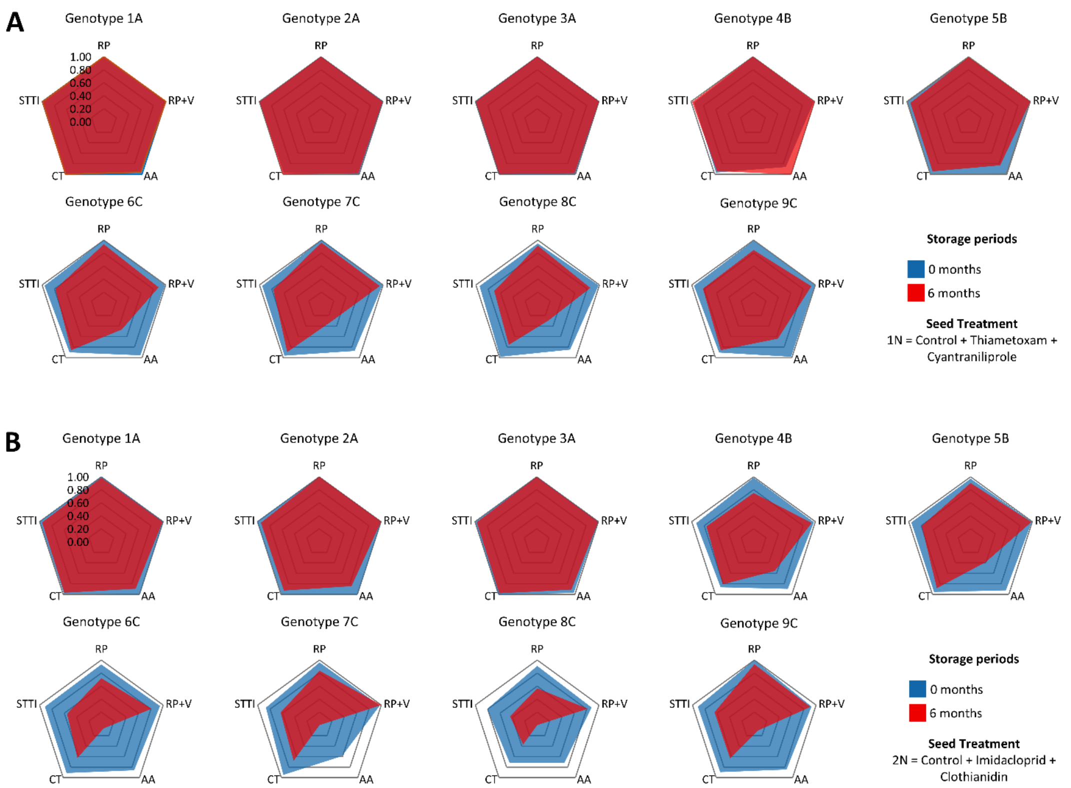


**Figure S1.** Radar plots illustrating the Seed Treatment Tolerance Index (STTI) for nine maize (*Zea mays* L.) hybrid genotypes across four physiological quality tests at time zero (blue area) and after six months of storage at 25 °C (red area). (A) Genotypes subjected to treatment 1N (control + one neonicotinoid insecticide + one diamide insecticide). (B) Genotypes subjected to treatment 2N (control + two neonicotinoid insecticides). Axes represent the STTI calculated for each test: RP (rolled paper germination), RP+V (rolled paper plus vermiculite germination), AA (accelerated aging), and CT (cold test). STTI values near 1.0 indicate full tolerance; values near 0 indicate complete sensitivity.

Supporting Information for: Potential methodology for phenotyping maize seed tolerance to storage after seed treatment (Reis et al., XXXX)

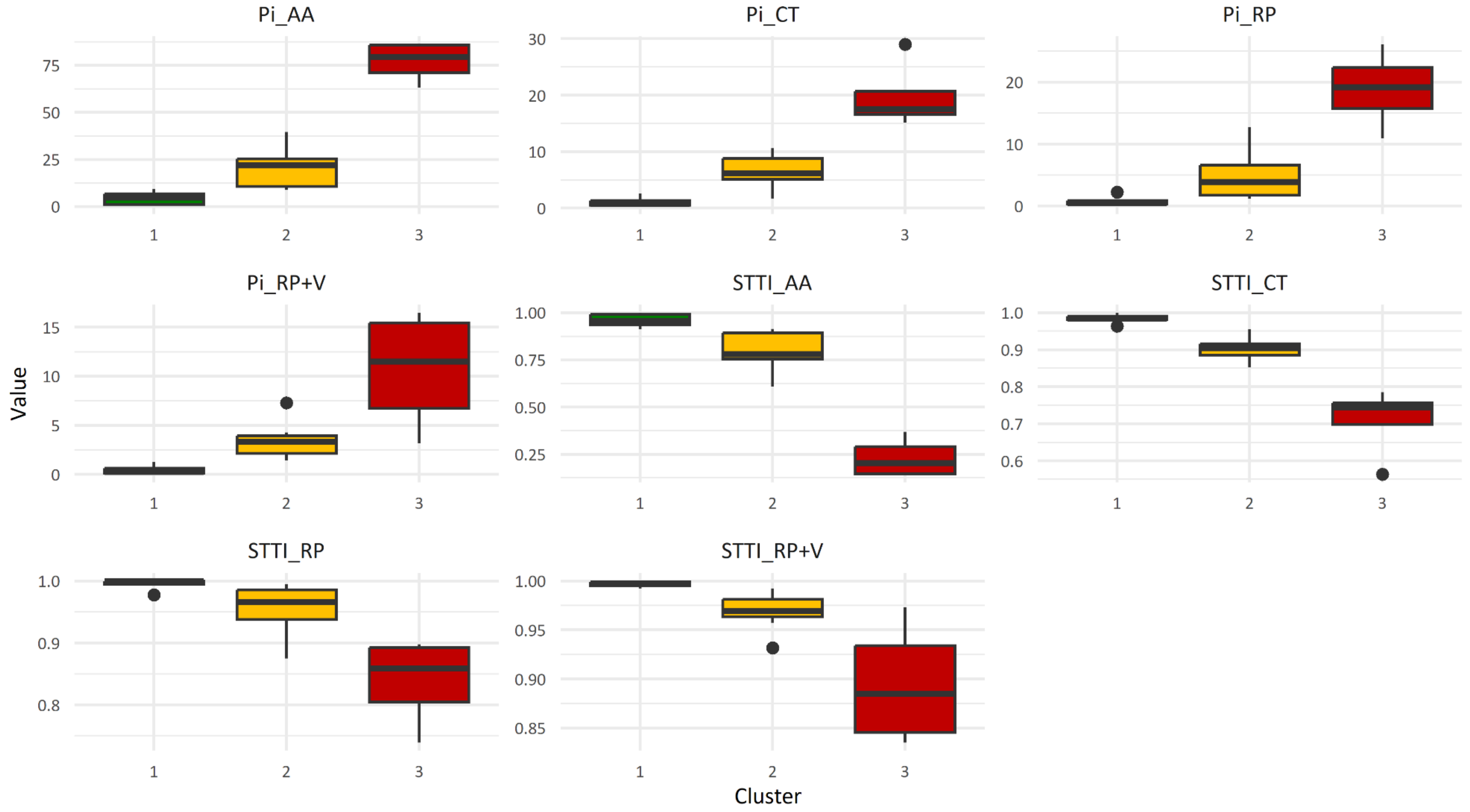


**Figure S2.** Boxplot comparison of the Phytotoxicity Index (Pi) and Seed Treatment Tolerance Index (STTI) across three statistically distinct genotype clusters (Groups 1, 2, and 3) identified through hierarchical cluster analysis and validated by MANOVA. Each panel represents an index calculated for a specific physiological test: AA (accelerated aging), CT (cold test), RP (rolled paper germination), and RP+V (rolled paper plus vermiculite germination). Different letters above boxplots indicate significant differences among groups by ANOVA (Scott–Knott test, $p \leq 0.05$).







Velikova, V., I. Yordanov, and A. Edreva. 2000. Oxidative stress and some antioxidant systems in acid rain-treated bean plants. Plant Science 151(1): 59–66. doi: 10.1016/S0168-9452(99)00197-1.

Ward, J.H. 1963. Hierarchical Grouping to Optimize an Objective Function. Journal of the American Statistical Association 58(301): 236–244. doi: 10.1080/01621459.1963.10500845.

Wilks, S.S. 1932. Certain Generalizations in the Analysis of Variance. Biometrika 24(3/4): 471. doi: 10.2307/2331979.
